# Screening, Isolation and Characterization of dye degrading bacteria from textile dye effluents

**DOI:** 10.1101/2021.12.20.473465

**Authors:** Shruti B. Sakpal, Kishori S. Tarfe

## Abstract

Textile dye industry waste is one among the foremost serious issues within the atmosphere. The dye wastes are severely harmful to surface water bodies. The dye degradation and decolorisation processes, that embody several physical and chemical strategies having inherent drawbacks, like cost accounting, economically impracticable (require additional energy and chemicals), unable to get rid of a number of the recalcitrant dyes and production of huge quantity of sludge that if not properly treated, successively will cause secondary pollution. So, biological degradation, being eco-friendly and cheap methodology, is taken into account as an efficient methodology for the removal of nephrotoxic radical dyes. Our present study was therefore aimed to isolate dyestuff decolorizing microorganism from dyeing industry effluent associate degreed to check their characteristics so as to use them as an economical bio agent for decolorizing and mineralizing nephrotoxic radical dyes.Various microorganism like *Bacillus subtilis, Aeromonas hydrophila* and *Bacillus Cereus*, fungi & actinomycetes are found to possess dye decolorizing activity. For the aim of finding out their characteristics, water sample was subjected to enrichment culture technique and then isolated on sterile nutrient agar plates containing 0.005%, 0.01%, and 1% of Congo red dye. The probable isolated organism from Congo red dye i.e. *Pantoea agglomerans* was found which can possess the ability to decolorize Congo red at lower concentration. The probable isolates obtained must be additional investigated relating to varied factors like dye degradation capability, media composition affecting dye degradation & mechanism of dye degrading activity.

## 1. INTRODUCTION

In modern life, rapid industrialization and urbanization resulted in the discharge of large amount of waste in to the environment, which in turn creates pollution. Water is essential for survival and existence of life on planet earth. The waste water and sewage are released from the industries, that wastes are entering into the water bodies, it is one of major source of environment toxicity [1], it also affect the soil micro flora and aquatic ecosystem [2]. The most environmental problem faced due to the textile dyeing industry is that the industry produces large volumes of high strength of aqueous waste effluents.

The discharge of dye effluents containing recalcitrant residue into rivers and lakes [3]. The residual dyes from different source such as textile industries, cosmetics, paper mills, pulp industries, dyeing and dye intermediates and bleaching industries, more than 80,000 tons of dyes and pigments are produced in these industries. Especially in textile industries produced more than 70% of the total quantity of waste in India [4]. India is the second largest exporter of dyestuffs and intermediates after China. The textile industry accounts for the largest consumption of dyestuffs, at nearly 80%.

The effluent which is untreated is one of the major sources of consumed metal dyes, phenol, aromatic amines [4, 5, 6]; several aromatic amines are known mutagens and carcinogens to human beings. Dyes also affect internal organ like kidney, liver, gastrointestinal tract. In ancient age natural sources were used for dying clothes. But the extraction process was a bit difficult and also expensive. Hence there was a need of synthetic dyes. In 1856, English chemist William Henry Perkin, in his experiment with aniline (one of the simplest chemical components of coal tar) obtained a black precipitate and discovered purple color, which readily dyed silk and was much more stable in sunlight than any other (natural) purple dye then in use. [7]

Dyes are natural or synthetic colored organic compounds having the property of imparting their color to the other substances, such as textile fibers. [7] Synthetic dyes are used extensively for textile dyeing, paper printing, leather dyeing, color photography and as additives in petroleum products because of their ease and cost effectiveness in synthesis, firmness, high stability to light, temperature, detergent and microbial attack and variety in color as compared to natural dyes [8] Approximately, 10,000 different dyes and pigments are used in different industries and their production exceeds over 7 × 10^5^ tons annually worldwide [9]

“Bioremediation” has become a key microbial tool to deal with different pollutants, is a key research area in the field of environmental science. [11] A number of bacteria, fungi, yeasts, algae and actinomycetes have been capable of decolorizing a range of azo dyes.[12] A number of biological and physico-chemical methods have been developed for the efficient removal of industrial azo dyes. Particularly, bacteria are the most frequently applied microorganisms for degradation of azo dyes, as they are generally fast to multiply rapidly under aerobic, anaerobic, facultative conditions as well as in extreme environmental conditions, like high salinity and wide variations in both pH and temperature[13] Microbes can concentrate, accumulate and absorb heavy metals inside cell or cell walls. Various microorganisms including, yeasts, *Proteus sp*., *Enterococcus sp*., *Streptococcus sp*., *Bacillus subtillis* and *Streptococcus sp*. have been previously isolated to degrade azo compounds [14]. Immobilized microorganisms are also being used for water purification e.g. immobilized mycelium of Coriolus versicolor is being used for removing colors/pigments from Kraft mill wastes.

Various bacterial strains reduce azo dyes under both anaerobic and aerobic conditions to the corresponding amines [10]. The bacteria *Sphingomonas xenophaga* BN6, *Agrobacterium tumefaciens*, *Ralstonia eutropha* 335, *Hydrogenophaga palleronii*, *Escherichia coli* K12 and *Flexibacter filiformis* (Gram negative), *Bacillus subtilis*, *Rhodococcus erythropolis* and *Lactobacillus plantarum*) (Gram negative) and Archea (*Halobacterium salinarum*) are reported to reduce azo dyes under anaerobic condition. [7] The decolorization of azo dyes by some bacterial strains of *Pseudomonas aeruginosa*, *Alcaligenes faecalis*, *Proteus mirabilis*, *Serratia marcescens* and *Bacillus licheniformis* at static and shaking conditions. *Pseudomonas aeruginosa* was able to detoxify the dye, Direct Orange 39(1,000 ppm each day) effectively. [15]

Attempts to isolate pure bacterial cultures capable of degrading azo dyes started way back in 1970s with isolation of *Bacillus subtilis*, *Aeromonas hydrophila* and *Bacillus cereus* [16]. Application of single bacterial cultures like *Proteus mirabilis*, *Pseudomonas luteola*, and *Pseudomonas sp*., has shown very promising results for the azo dye degradation under anoxic conditions [12].Other bacterial strains of *Desulfovibrio desulfuricans, Exiguobacterium sp., Sphingomonas sp., Rhizobium radiobacter* and *Comamonas sp*. were also reported for decolorization of various azo dyes. Among these strains, *Pseudomonas* is widely used for decolorization study of azo dyes and also exploited widely to decolorize commercial textile azo dyes, such as Red BLI, Reactive Red 2, Red HE7B, Reactive Blue 172, Reactive Red 22, Orange I and II. In addition, Micrococcus sp. is an interesting example, which decolorizes dyes faster under aerobic conditions than in anaerobic environments. [11]

Bacteria capable of dye decolorization/biodegradation either in pure cultures or in consortia have been reported [11, 14-17]. However, comprehensive solutions for sulfonated azo dyes removal are far from reality, which calls for continued search for new organisms and technologies.

This study aimed to isolate and characterize an efficient bacterial strain, which exhibited the remarkable ability to degrade Congo red, methylene blue and malachite green dye from which, results were obtained only for Congo red dye. Therefore further study was carried out by using Congo red dye only.

## 2. MATERIALS AND METHODS

The study was conducted at Department of Biotechnology, Smt. Chandibai Himathmal Mansukhani College, Ulhasnagar-3, District Thane, Maharashtra, India.

### 2.1 Sample Collection

Contaminated water samples collected from Ulhas River, Ulhasnagar east were used as a source to isolate bacteria with distinct morphological colony characters capable of decolorizing selected azo dye. The water samples were collected in airtight bottles and filtered through ordinary filter paper to remove large suspended particles and the filtrate were used for the isolation procedure.

### 2.2 Physico-chemical property analysis

The collected effluent samples have been analyzed to determine its physico-chemical parameters. The various parameters like Chemical oxygen demand (COD), Biological oxygen demand (BOD), Dissolved Oxygen (DO) were analyzed in the laboratory by the standard protocol.

### 2.3 Enrichment and isolation of dye tolerating strains-

#### a) Enrichment

The water sample collected were subjected to enrichment culture technique.

The enrichment was carried out in 100 ml Nutrient Broth medium containing 1% Congo red taken in 250 ml Erlenmeyer flask by adding 10 ml of sample water. The flask was then incubated in a rotary shaker at 50 rpm at 30°C for one week.

#### b) Isolation and Characterization of dye degrading bacteria

A loopful from enriched flask was streaked on media plate viz. sterile Nutrient agar with 1% dye concentration Congo red and was incubated at room temperature for 24 hours. Similarly, effluent sample was directly streaked on sterile nutrient agar plates with 0.005% and 0.01% and 1% Congo red and incubated at room temperature for 24 hours. The identification of dye degrading bacterial strains was carried out based on morphological characteristics of colonies obtained on plates.

#### c) Staining and motility

24 hr. old cultures of all the isolates were used to study Gram reaction and the cell morphology. The morphological characterization of the isolates was checked by Gram’s staining.

#### d) Growth curve

20 ml of sterile nutrient broth was inoculated with 18 hrs. Old culture of 0.005%, 0.01%, 1% concentration of Congo red dye in side arm flask. The maximum absorption was noted at 545nm after every 30 minutes. Optical density was noted and a graph was plotted denoting x-axis – time in minutes and y-axis – optical density at 545nm.

## 3. OBSERVATION AND RESULTS

Textile effluents are directly released into Ulhas River at Ulhasnagar. Thus, only one effluent sample from Ulhas River, Ulhasnagar East was used.

### 3.1 Physico-chemical property analysis

#### i) Biological oxygen demand

1:100 diluted sewage sample.

|  |  |  |
| --- | --- | --- |
|  | D.O.<br>(mg/liter) | B.O.D.<br>(mg/liter) |
| Day- 1 | 170 | 80 |
| Day- 5 | 90 |  |

#### ii) Chemical oxygen demand

Normality (N) = 0.1, Volume of sample = 10 ml

|  |  |  |
| --- | --- | --- |
| Blank reading | Test Reading | C.O.D. |
| 2.3 ml | 0.9 ml | 112 mg/liter |

### 3.2 Colony characteristics of selected strains

Plates containing 0.005%, 0.01% and 1% of Congo red dye with sterile Nutrient agar medium showed growth of colonies which was of 1mm size, circular shape, reddish white color, raised elevation, mucoid consistency, opaque and gram negative which indicated that the bacteria belongs to Coccobacilli genera.

**Figure 1:**
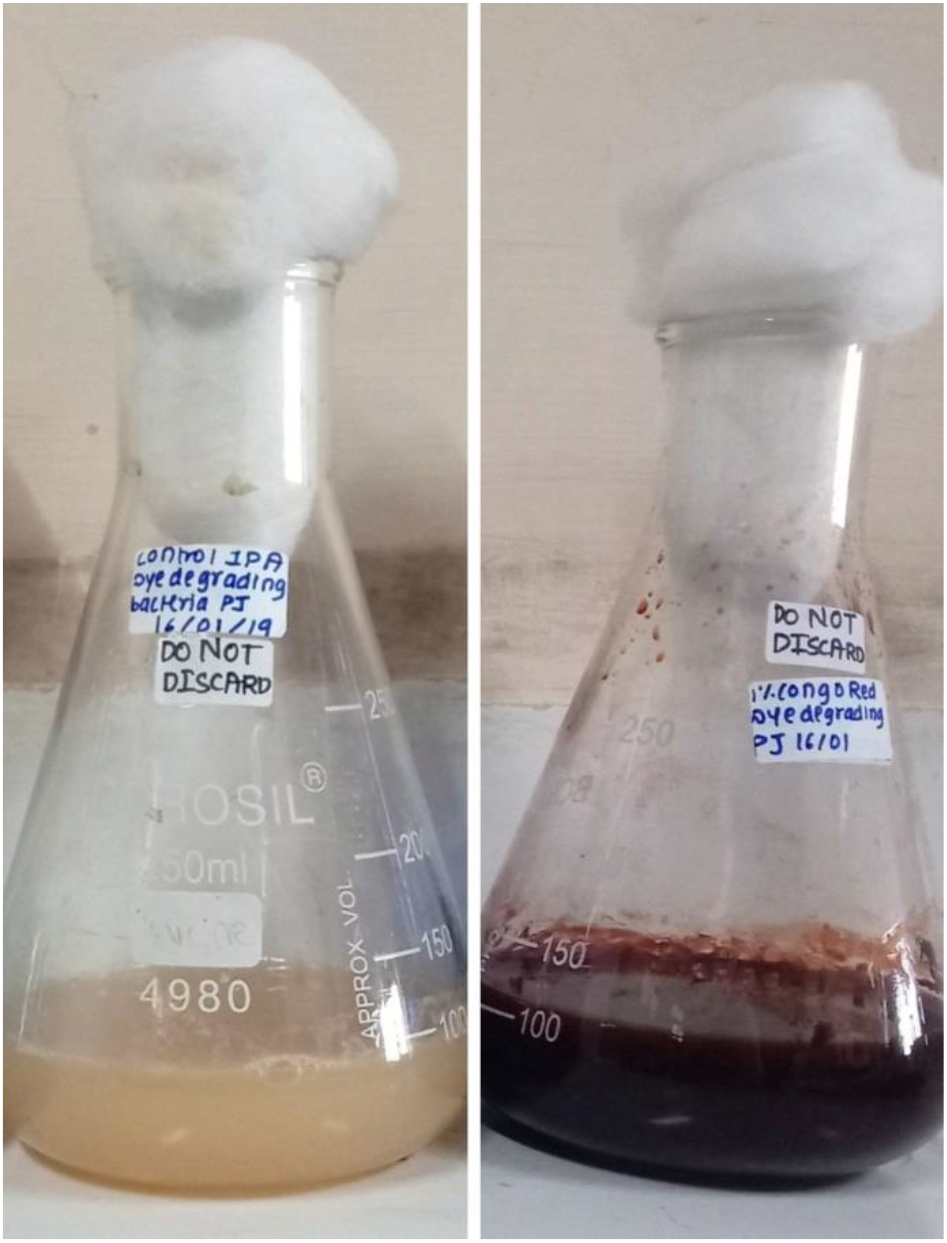
Enrichment of dye degrading bacteria from textile of effluent sample.

**Figure 2:**
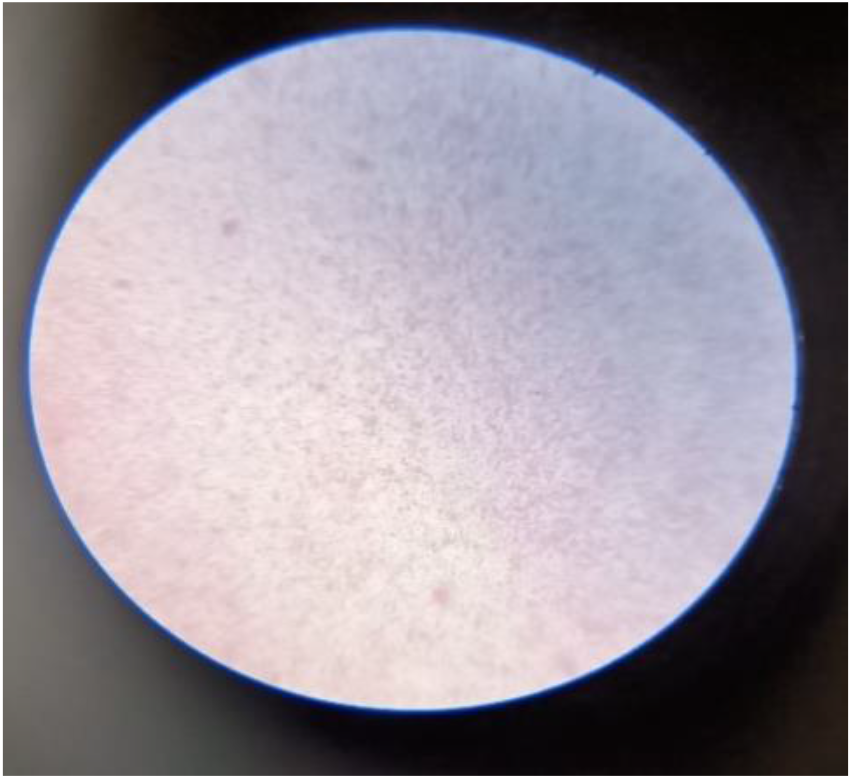
Bacteria isolated from Congo Red plate.

### 3.3 Growth Curve Analysis

Absorbance at 545nm

**Figure 3:**
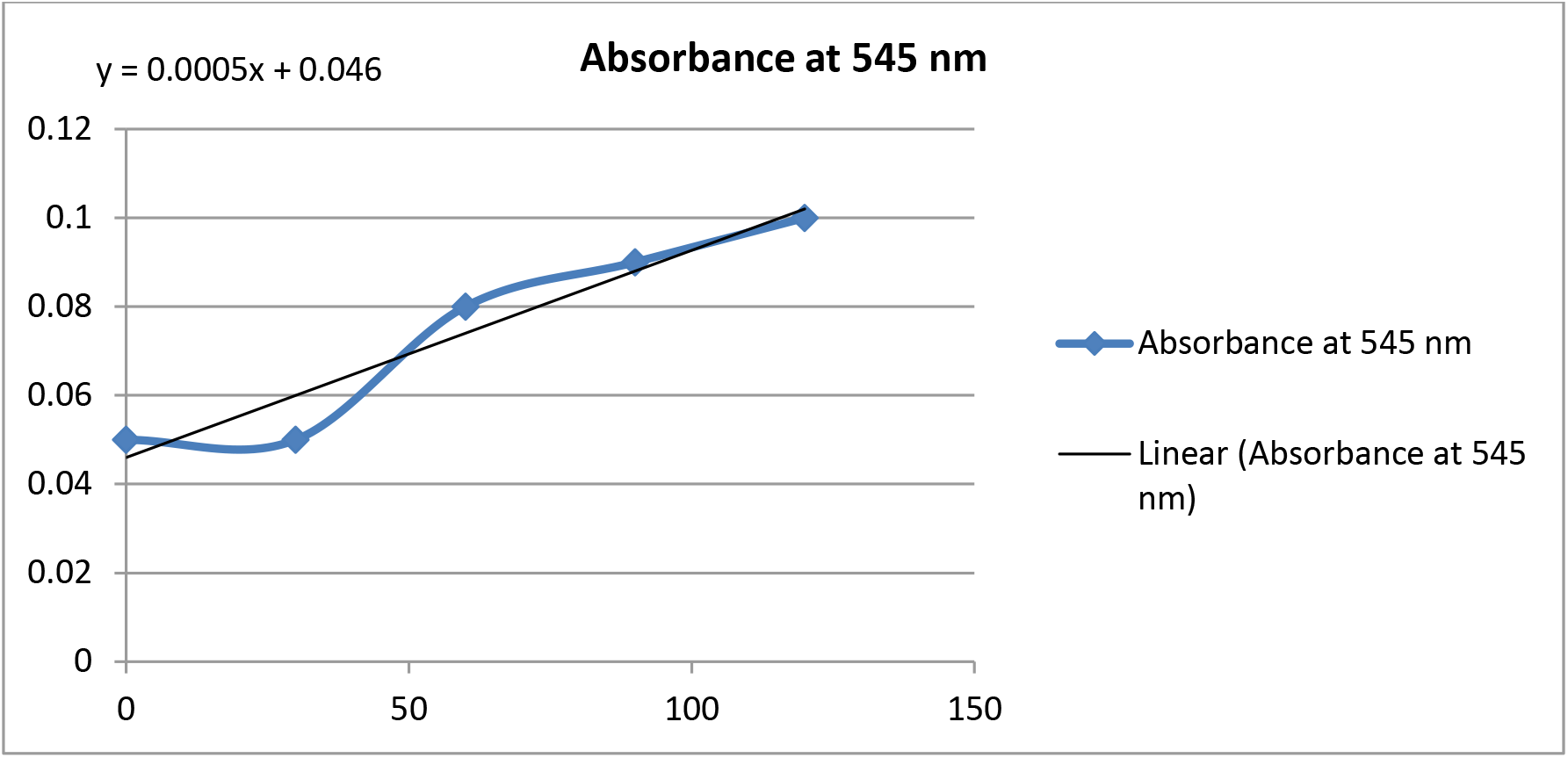
Growth curve for bacteria grown in 0.005% of Congo Red.

**Figure 4:**
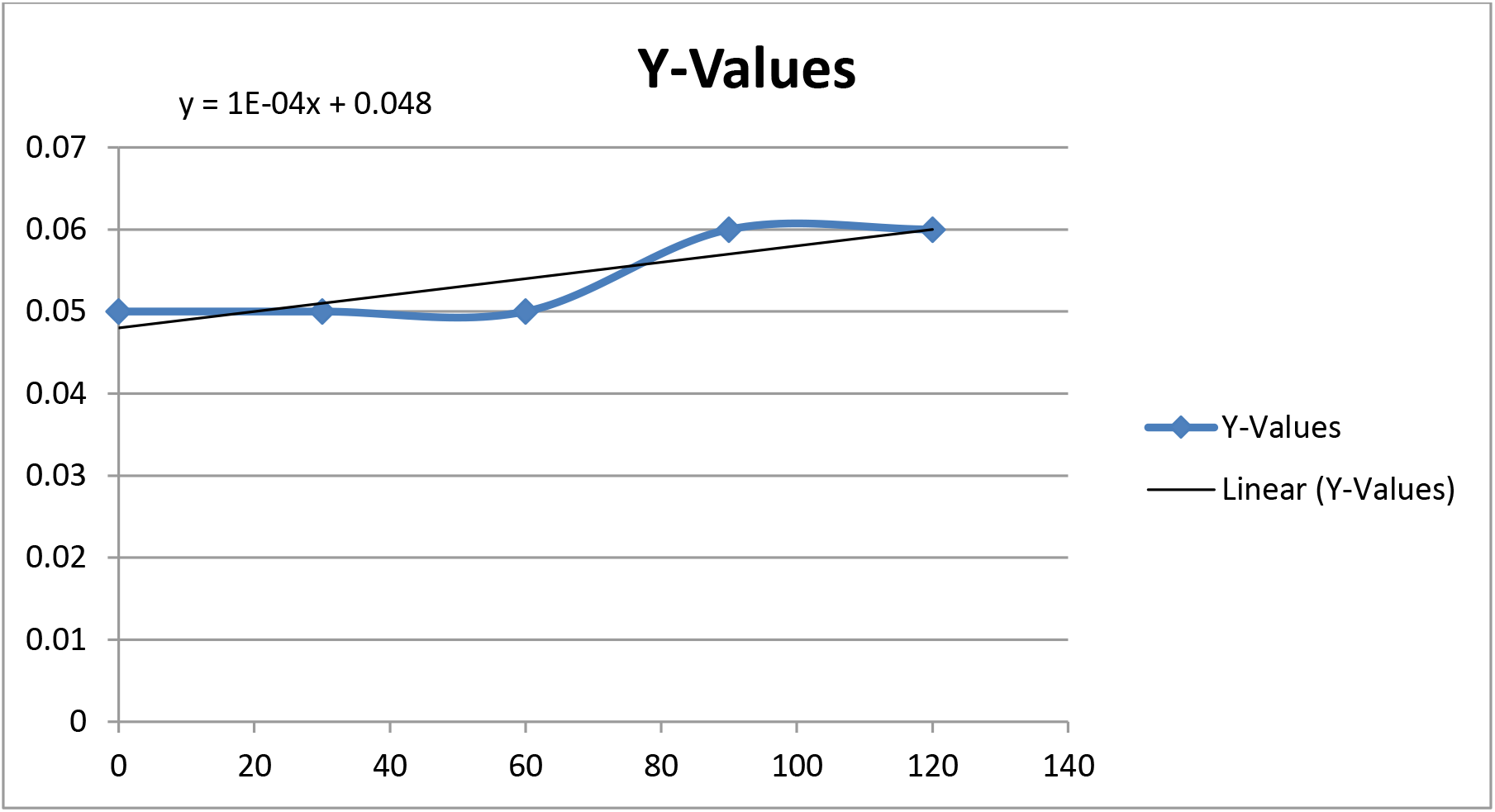
Growth curve for bacteria grown in 0.01% of Congo Red.

**Figure 5:**
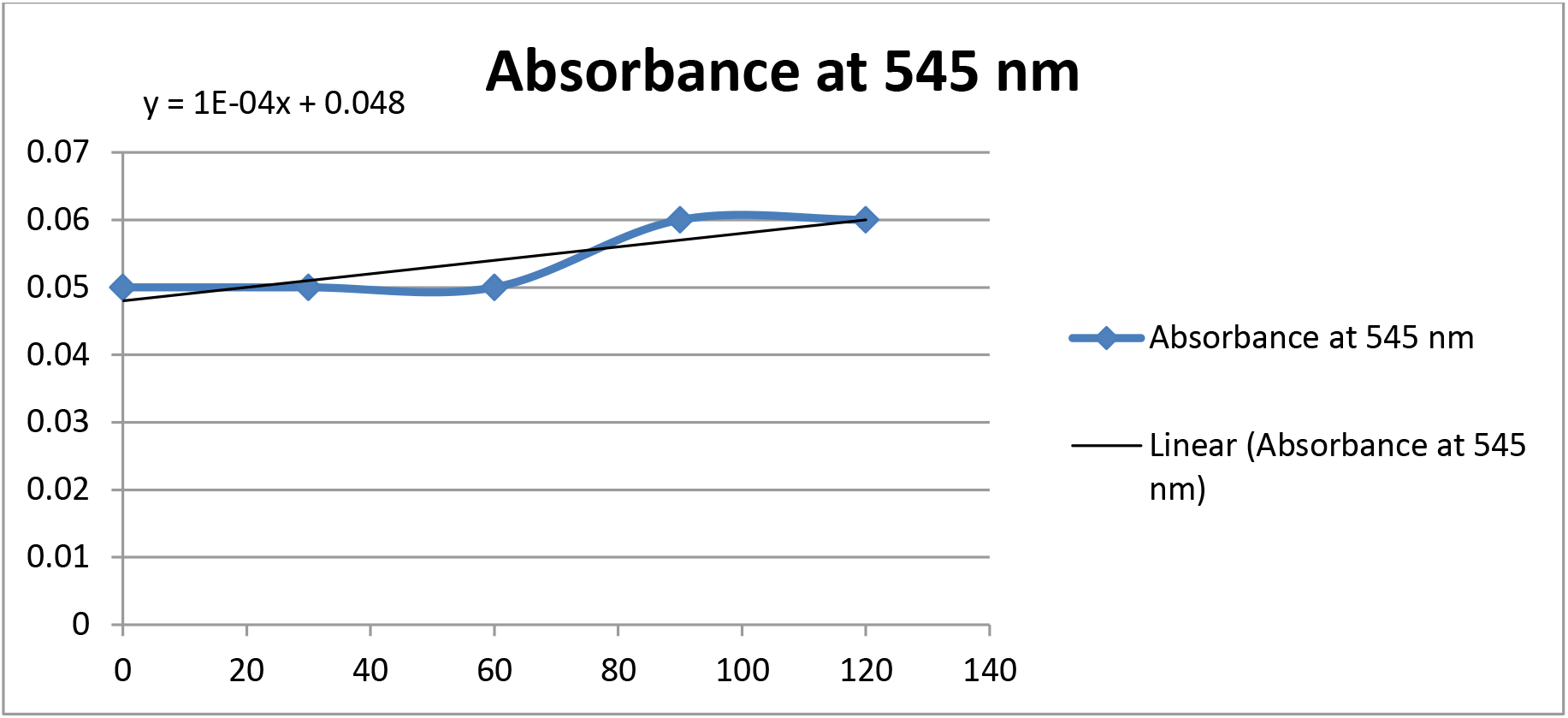
Growth curve for bacteria grown in 1% of Congo Red.

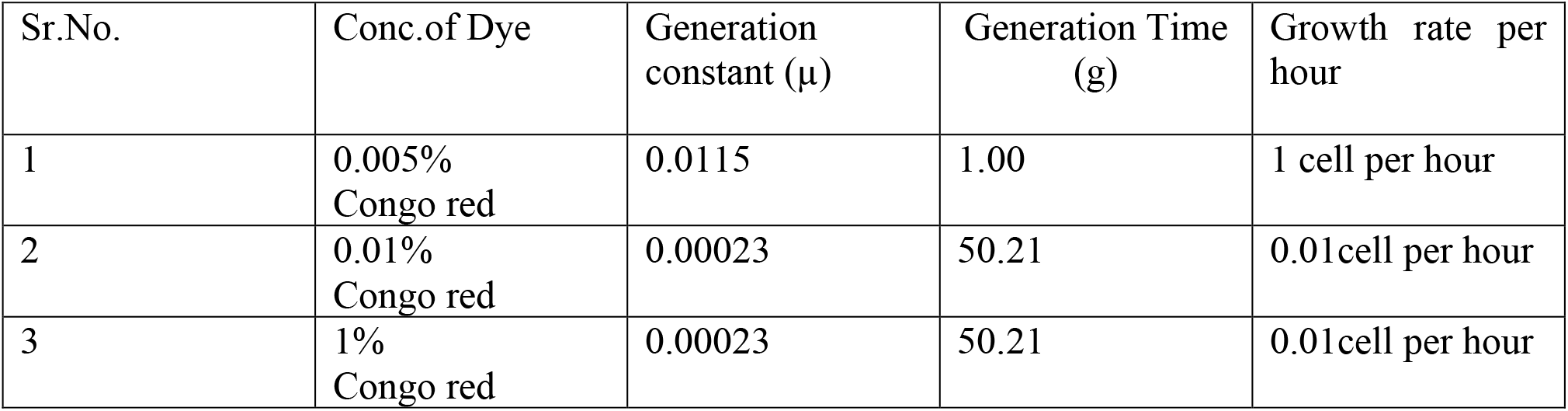

**Figure 5: Growth curve for bacteria grown in 1% of Congo Red.**
| Sr.No. | Conc.of Dye | Generation constant (μ) | Generation Time (g) | Growth rate per hour |
| --- | --- | --- | --- | --- |
| 1 | 0.005% Congo red | 0.0115 | 1.00 | 1 cell per hour |
| 2 | 0.01% Congo red | 0.00023 | 50.21 | 0.01cell per hour |
| 3 | 1% Congo red | 0.00023 | 50.21 | 0.01cell per hour |

## 4. DISCCUSION

Textile dye industrial effluents are one in all major sources of environmental toxicity. It not solely affects the standard of water however conjointly has injurious impact on the microflora and aquatic ecosystems. The dye effluents were collected from Ulhas river in Ulhasnagar. The industries located in Ulhasnagar discharge the various colored effluents with dyes and deadly compounds into the open setting. It was found that the dyeing industry is among those that contribute to pollution. Therefore, the collected sample were analyzed to determine their physico-chemical characteristics of the dye effluents like B.O.D.,COD and D.O. content of effluent water.

The B.O.D. and COD of effluent sample was found to be 80 mg / liter and 112 mg / liter severally. The B.O.D. values was found to be higher than the permeasible values. Enrichment and isolation of bacterium from effluent sample was conjointly done in sterile nutrient agar media with different concentration of Congo red dye such 1%, 0.01% and 0.005%. After isolation, Gram staining of isolated bacterium on Congo red plates was performed to identify the bacteria and it had been found that the isolated bacteria could be a Gram negative bacterium. Growth curve methodology was performed to find out the growth rate of bacteria per hour. It was found that 1 cell per hour, 0.01 cell per hour and 0.01 cell per hour were the growth rates of bacteria in a media containing 0.005%, 0.01% and 1% of Congo red dye, respectively. Dye degrading bacterium known as *Pantoea agglomerans*.

The bacterial isolates are often screen for the decolourization of dye effluents. The organisms isolated must be further investigated regarding numerous factors like media composition affecting dye degradation and mechanism of dye degrading activity. Immobilized cells are often utilized in bioreactor in Textile industries as a treatment of dye effluent.Study conducted by Alyssa M. Walterson and John Stavrinides reveals that bacterial genus Pantoea was discovered about 25 years ago but approximately 20 species are yet known. Isolates that are been obtained from water and soil are mostly used in biodegradation and bioremediation of toxic products released from industries into the environment.

## 5. CONCLUSION

In this Investigation, enrichment of dye degrading bacteria in a nutrient agar containing different concentrations of Congo red dye was carried out. After enrichment for 24 hrs. loopful culture was streaked on media plates and incubated for 24 hrs. at R.T. to isolate dye degrading bacterium. Well isolated colonies were found which then studied for their characteristics like cell shape, size, colour, arrangement, etc. By performing Gram’s Staining, the Gram nature of all the isolated organisms was found to be Gram negative coccobacilli. The probable organism isolated that decolorizes the Congo red was can be *Pantoea agglomerans*.

As this can be an issue of environmental pollution, the isolates will play a crucial role in the prevention of pollution. It can be concluded that the dye decolorizers are often can be for the treatment of waste water and textile effluents that are alarming for the environment. At future studies, the characterization, optimization and molecular investigation of the dyestuff degradation by the isolate also can provide important info during this field to solve the arising problem. We will able to conjointly develop molecular biology technique and commercially can produce such enzyme to protect environment from the pollution.

## 6. ACKNOWLEDGEMENT

Authors are thankful to Department of Biotechnology, Smt. Chandibai Himathmal Mansukhani College, Ulhasnagar, District Thane, Maharashtra, India.

## REFERENCES

[1] Arminder Kaur, Siddharth Vats, Sumit Rekhi, Ankit Bhardwaj, Jharna Goel, Ranjeet S. Tanwar and Komal. K. Gaur, Procedia Environmental Sciences, 2010, 2, 595–599.

[2] Thoker Farook Ahmed, Manderia Sushil1 and Manderia Krishna, International Research Journal of Environment Sciences, 2012, 1(2), 41–45.

[3] N. Manikandan, S. SurumbarKuzhali and R. Kumuthakalavalli, J. Microbiol. Biotech. Res., 2012, 2(1), 57–62.

[4] K. Rajeswari, R. Subashkumar and K. Vijayaraman, J. Microbiol. Biotech. Res., 2013, 3 (5), 37–41.

[5] K. Varunprasath and A.N. Daniel, Iranica. J. Energy Environ., 2010, 1, 315–320.

[6] D. Suteu, C. Zaharia, D. Bibla, A. Muresan, R. Muresan and A. Popescu, Industria Textila, 2009, 5, 254–263.

[7] Lokendra Singh and Ved Pal Singh Textile Dyes Degradation: A MicrobialApproach for Biodegradation of Pollutants, Research gate DOI: 10.1007/978-3-319-10942-8_9

[8] Couto SR (2009) Dye removal by immobilised fungi. Biotech Adv 27:227–233

[9] Zollinger H (1987) Color chemistry synthesis, property and application of organic dyes and pigments. VCH publishers, New York, pp 92–102.

[10] Standard Methods for the Examination of Water and Wastewater, American Public Health Association, American Water Works Association, Water Environment Federation.

[11] Shailesh R. Dave, Tallika L. Patel and Devayani R. Tipre (2015) Application of novel consortium TSR for treatment of industrial dye manufacturing effluent with concurrent removal of ADMI, COD, heavy metals and toxicity.

[12] Rijuta Ganesh Saratale, Jo-Shu Chang, Ganesh D. Saratale, Sanjay P Govindwar Bacterial Decolorization and Degradation of Azo Dyes: A Review, Journal of the Taiwan Institute of Chemical Engineers January 2011.

[13] Solis M, Solis A, Perez HI, Manjarrez N, Flores M (2012) Microbial decoloration of azo dyes: a review. Process Biochem 47:1723–1748

[14] Brown JP (1981) Reduction of polymeric azo and nitro dyes by intestinal bacteria. Appl Environ Microbiol 41(5): 1283–1286.

[15] Jadhav JP, Phugare SS, Dhanve RS, Jadhav SB (2010) Rapid biodegradation and decolorization of Direct Orange 39 (Orange TGLL) by an isolated bacterium Pseudomonas aeruginosa strain BCH. Biodegradation 21:453–463

[16] Wuhrmann K, Mechsner K, Kappeler T (1980) Investigations on rate-determining factors in the microbial reduction of azo dyes.

